# Turning mechanics of the bipedal hopping Desert kangaroo rat (*Dipodomys deserti*)

**DOI:** 10.1101/509398

## Abstract

A fundamental aspect of animal ecology is the ability to avoid getting eaten. Catching prey and avoiding depredation involves a dynamic interplay between forward and lateral acceleration. Success at these tasks depends on achieving sufficient performance. In turn, performance is determined by biomechanics. After observing several kangaroo rats utilize turns under the duress of simulated predation in the field, we designed and conducted an experiment in the lab to measure turning mechanics of desert kangaroo rats. The average turning speed in our study was 1.2 ms^-1^. While field performances are rarely replicable in the lab, we found that kangaroo rats utilize braking impulses (~0.4 N), followed by lateral impulses, and orientated their body early in the turn to match trajectory change. to execute a turn. Coordinating turn in this ways likely prioritizes safety.

## INTRODUCTION

Locomotion is fundamental to many animal behaviors (Dickinson *et al.* 2000). Locomotor performance *(e.g.* speed) in terrestrial vertebrates depends on multiple, interacting intrinsic and extrinsic factors relative to the musculoskeletal system. Extrinsic examples limiting locomotor performance include the mechanical properties of the terrain (Bergman *et al.* 2017; Collins *et al.* 2013; Lejeune and Willems 1998), temperature (Swoap *et al.* 1993; Jayne *et al.* 1990), and ecological context (Wilson *et al.* 2015). Intrinsic factors include the ability to apply an impulse to Earth’s surface via muscular contraction (Biewener 2016). Understanding the mechanics underpinning locomotion are necessary for understanding how structures evolved to meet the functional demands of behaving in challenging environments.

Under the duress of potential predation, sprint speed may enhance survival in many animals (Husak 2006 a&b; Miles 2006). As a result, many laboratory-based experiments quantify straight-forward sprints. These studies illuminate the relationships between morphology, mechanics, and linear progression. Yet, field studies and mathematical modeling reveal a vast repertoire of locomotor behaviors are employed in natural conditions. In fact, the ability to change direction during a predator-prey pursuit often means the difference between escape success and capture (Moore *et al.* 2015; Wilson et al. 2013 a&b).

Understanding how species evolved biomechanical traits enabling larger impulses and faster limb cycling have undergone extensive investigation in many vertebrates. Terrestrial sprint speed is limited by how quickly limbs swing and how much force is applied to the ground (Blickhan 1989; Cavagna 1976; Usherwood and Wilson 2005; Weyand 2000). Applying force to the ground over a period of time results in an impulse, causing the mass of the animal to increase or decrease velocity. Greater and more rapid force production during stance can produce higher net impulse, thereby increasing speed. The timing, magnitude, sequence, and direction of forces applied to the ground as measured under experimental conditions represent how natural selection has acted on the coordination and tradeoffs of a given locomotor behavior (Collins and Higham 2018; Irschick and Garland 2001; McElroy *et al.* 2012).

Turning by terrestrial animals consists of decelerating the center of mass in an initial trajectory and accelerating in the trajectory of a new angle, as well as orienting the body to face the new trajectory (Figure 1). In other words, an individual changes its direction of movement, and looks in the new direction (Jindrich and Full 1999). Changing trajectory requires a force to be applied to the center of mass and therefore can only occur while contacting the ground. However, orientation change may occur on the ground or in aerial phase by moving or changing the shape of an appendage such as a tail via the law of conservation of momentum (Alexander 2002; Carrier *et al.* 2001; Lee *et al.* 2001). Turning animals will topple if lateral acceleration overcomes fore-aft acceleration (Alexander 2002). Lower centers of mass and wider bodies are morphological components that help prevent toppling and learning into a turn will ameliorate this danger (Alexander 2002).

**Figure 1:**
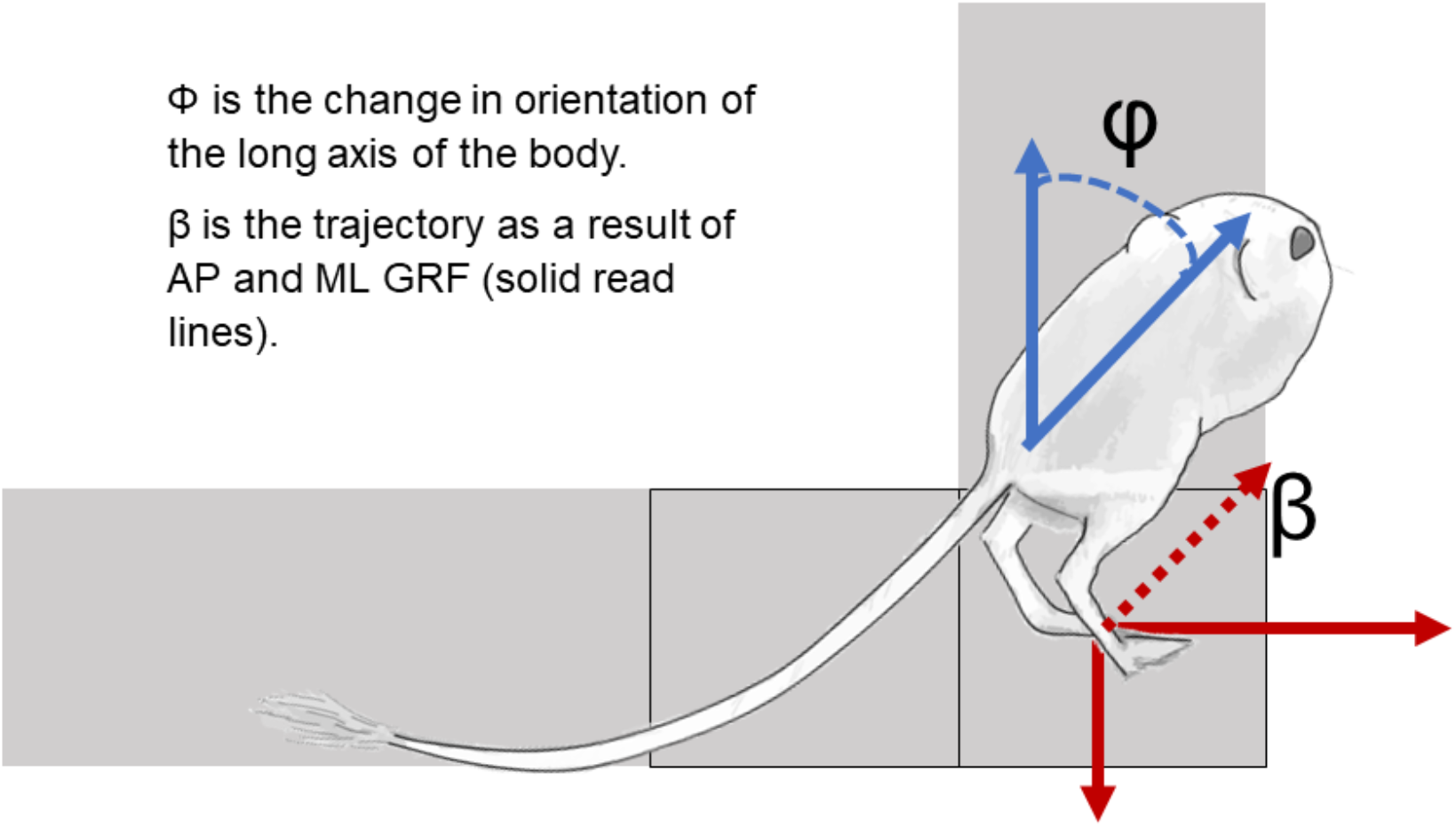
A schematic of the runway (grey), force plates (squares) and a hand-drawn *D. deserti,* illustrating change in orientation and trajectory.

Previous research on turning has focused on cutting and steering in running and walking humans, birds, and some quadrupedal animals (Carrier *et al.* 2001; Chang and Kram 2007; Demes 2006; Usherwood 2006 a&b). In humans, the horizontal forces necessary for turnings are produced by longer stances and shorter swings and running on a bend results in lower velocities (Greene and McMahon 1979; Usherwood and Wilson 2005). These results indicate that turning performance is limited by limb muscle force production. Furthermore, bipedally running humans exhibit functional decoupling of outside vs. inside limb mechanics, as force production appears to be most limited by the inside limb during bend track running (Chang and Kram 2007). Ostriches minimize changes in leg kinematics and changes to net torque production at their joints and utilize crossover and sidesteps during maneuvering (Jindrich *et a.l* 2007). Turning in quadrupeds varies by species, but many exhibit axial bending (Eliam 1994; Walter 2003). Substrate influences turning strategies in non-human primates and vary by species – whereas forces are more evenly distributed between front and hind limbs in Patas monkeys, lemurs exhibit hindlimb dominance in weight support (Demes 2006). Highly specialized, artificially selected greyhound dogs (and likely most cursorial quadrupeds) move by extending the back and torque about the hips while weight support occurs about the forelimbs. Spring-like muscular hip retractors working in conjunction with forelimbs enable greyhounds to power locomotion ‘virtually independent’ of supporting their mass. Therefore, the increase in effective weight when running through a turn does not cause decreased speed (Usherwood and Wilson 2005).

Bipedal hopping, in which the hindlimbs simultaneously contact the ground with no input from the forelimbs, is present in six extant mammal lineages (McGowan and Collins 2018). Bipedally hopping rodents utilize rapid direction changes and are putatively adapted to unpredictable locomotor trajectories as well as powerful accelerations (McGowan and Collins 2018). Their morphology is typically characterized by elongated and simplified distal elements of the hindlimb, reduced forelimbs, long tails, and specialized feet (McGowan and Collins 2018). Therefore, turning mechanics will likely differ from bipedal walkers and runners. After observing simulated predation attempts (Movie S1), we test the hypothesis that kangaroo rats *(Dipodomys deserti* Stephens 1887) change their trajectory by first applying a braking impulse and then a lateral impulse. If kangaroo rats first apply a braking force, then the lateral forces necessary to change trajectory are reduced. Second, we ask how changes in orientation are achieved and test whether they closely track changes in trajectory. Finally, we measure body lean to understand how this mechanism is used to ameliorate toppling danger. Our study reveals valuable insight into the evolution of bipedalism and maneuvering during hopping locomotion.

## MATERIALS AND METHODS

Twelve adult desert kangaroo rats, *D. deserti* (mass 84 – 110 g), were trapped from a long-term field site in Gold Butte National Monument near Mesquite, NV with permission from and under the guidance of the Nevada Department of Wildlife and the Bureau of Land Management. and returned to the campus of the University of Idaho (Moscow, Idaho) for experiments following a protocol approved by the Institutional Animal Care and Use Committee. Individuals were subject to quarantine for six weeks at the University of Idaho Animal Care Facility in which they were fed wild bird seed mix and spinach while under supervision of veterinary and animal care staff.

We constructed a custom racetrack measuring two meters long by 0.25 meters wide that was angled 90° at the center. The racetrack walls consisted of plexiglass to allow recording by three high-speed cameras (Xcitex Inc, Woburn, MA, USA) recording at 200 Hz. At the corner and adjacent to the corner were two force plates (AMTI HE 6×6, Watertown, MA, USA) recording at 600 Hz (Figure 1). The floor of the racetrack was constructed of painted wood covered by a PVC mat to enhance friction during steady and turning locomotion.

Prior to recording trials, a proxy for the center of mass (COM, near the ischium), and the base of the tail were marked using a combination of non-toxic white paint and black ink. These markers, plus the right or left eye, and the longest toe of the left or right foot were digitized in DLTdv5 (Hedrick 2008). The following were extracted from the digitized trials using a custom MATLAB script: COM trajectory and velocity, body orientation, body lean (the angle between the longest digitized toe and the COM), and anterior-posterior and medio-lateral forces. A low-passfourth-order Butterworth filter was applied with a cut-off frequency of 30 Hz was applied prior to analysis. We calculated the total change in trajectory and change in body orientation at three points in each trial. We analyzed the entry speed, the speed at the corner of the turn, and the exit speed, as well as the changes in trajectory and orientation corresponding to entry, corner, and exit. Vector changes were calculated by dividing the difference between 3D COM positions during each swing and stance and then calculating the angle of change between trajectories of each stride cycle by using the dot product.

Two to three individuals completed 20 – 50 runs per day in the racetrack for a total of 200 – 500 runs per individual. The individuals were encouraged to hop through the racetrack by pinching the tail, tapping on the dorsum, and clapping. Additionally, animals were allowed to hop uncoerced when possible. We observed no signs of exhaustion or stress before, during, or after trials. Recordings from the force plates and high-speed video were synchronized using a custom-built external trigger. We used VirtualDub and Windows Media Player to visually evaluate all trials upon completion of the experiment. Trials in which an animal failed to hop cleanly on both force plates and trials in which the animals failed to run “steadily” through the turn were discarded. The remaining trials included three to ten samples per individual. Six of the 12 *D. deserti* successfully hopped on the force plates in our custom racetrack. Therefore, we analyzed a reduced dataset.

We combine Spearman’s Rho and Pearson-product moments, to test our hypothesis of turning mechanisms. Data from multiple trials of all individuals are used to quantify the mechanics of turning. Because of the large amount of information and relatively small sample size, we used a significance cutoff < 0.02 and an R^2^ equal to or greater than 0.4. Significant results are presented in graphical format here. All analyses were performed in JMP© 12 (2016, SAS Institute). When pooled, all variables were normally distributed and relatively free of kurtosis and skew.

## RESULTS

In our study, left turns occurred more often than right turns, except for one individual that turned right nearly exclusively. Thus, our data describe entry and corner mechanics, but not exits. However, our results combined with those exits from a limited portion of trials indicate most of the changes in trajectory and orientation occur before or during the corner (~84°). Quadrupedal bounding was more common than bipedal hopping in our study (~90% quadrupedal). Differences between quadrupedal and bipedal turns are presented in Table 1. Generally, quadrupedal entries were faster than bipedal entries (1.47 ms^-1^ and 1.37 ms^-1^) but this difference was not statistically significant (p>0.05). Accordingly, trajectory, orientation, and impulses not different in quadrupedal turns relative to bipedal turns, but some of the forces were distributed between fore and hindlimbs during quadrupedal turns (Table 1). Quadrupedal turns often included stopping at the corner. Thus, many quadrupedal turns did not meet the criteria for inclusion in this study, leading to smaller than anticipated sample size (12 total quadrupedal turns, 24 bipedal turns). Overall trends in our experiment are that individuals exit the corner slower than entry, and that most of the turning occurs before or during the corner (Table 1). Representative force traces are presented in Figure 2.

**Figure 2:**
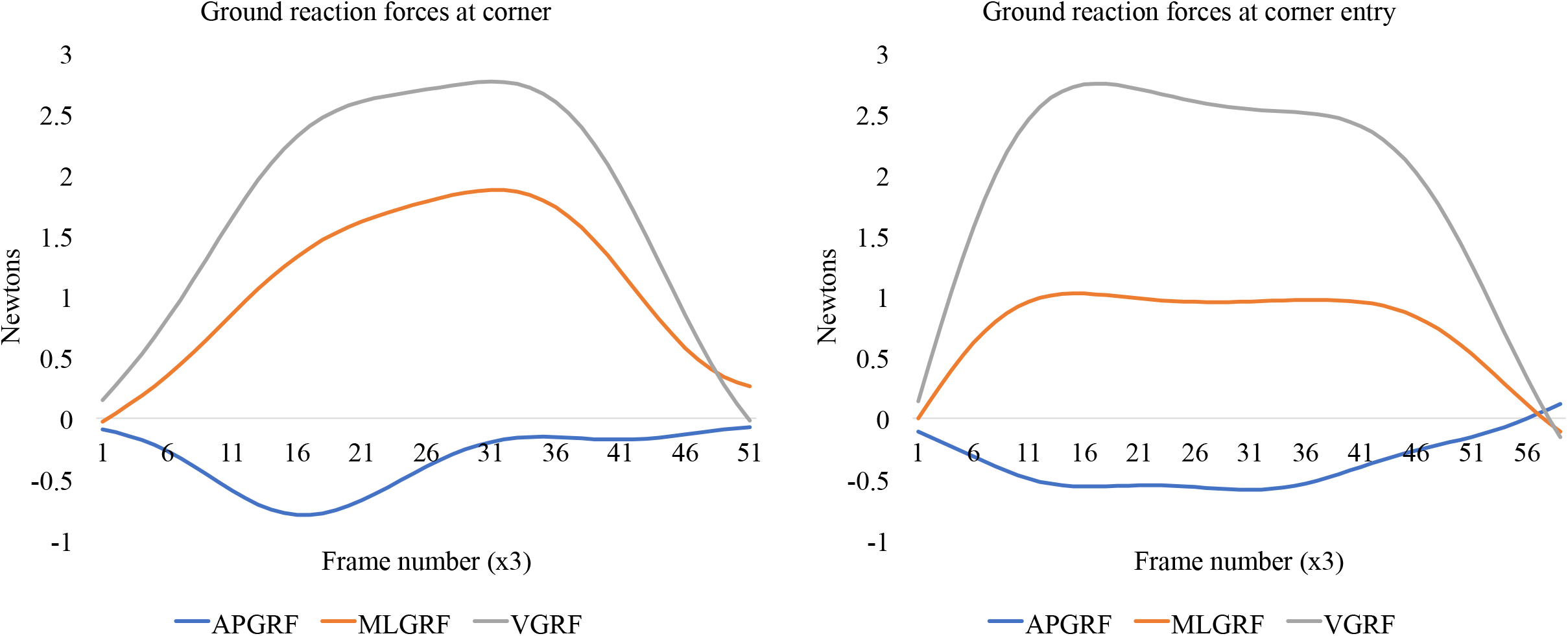
Example ground reaction force traces for the entry and corner force places.

**Table 1:**
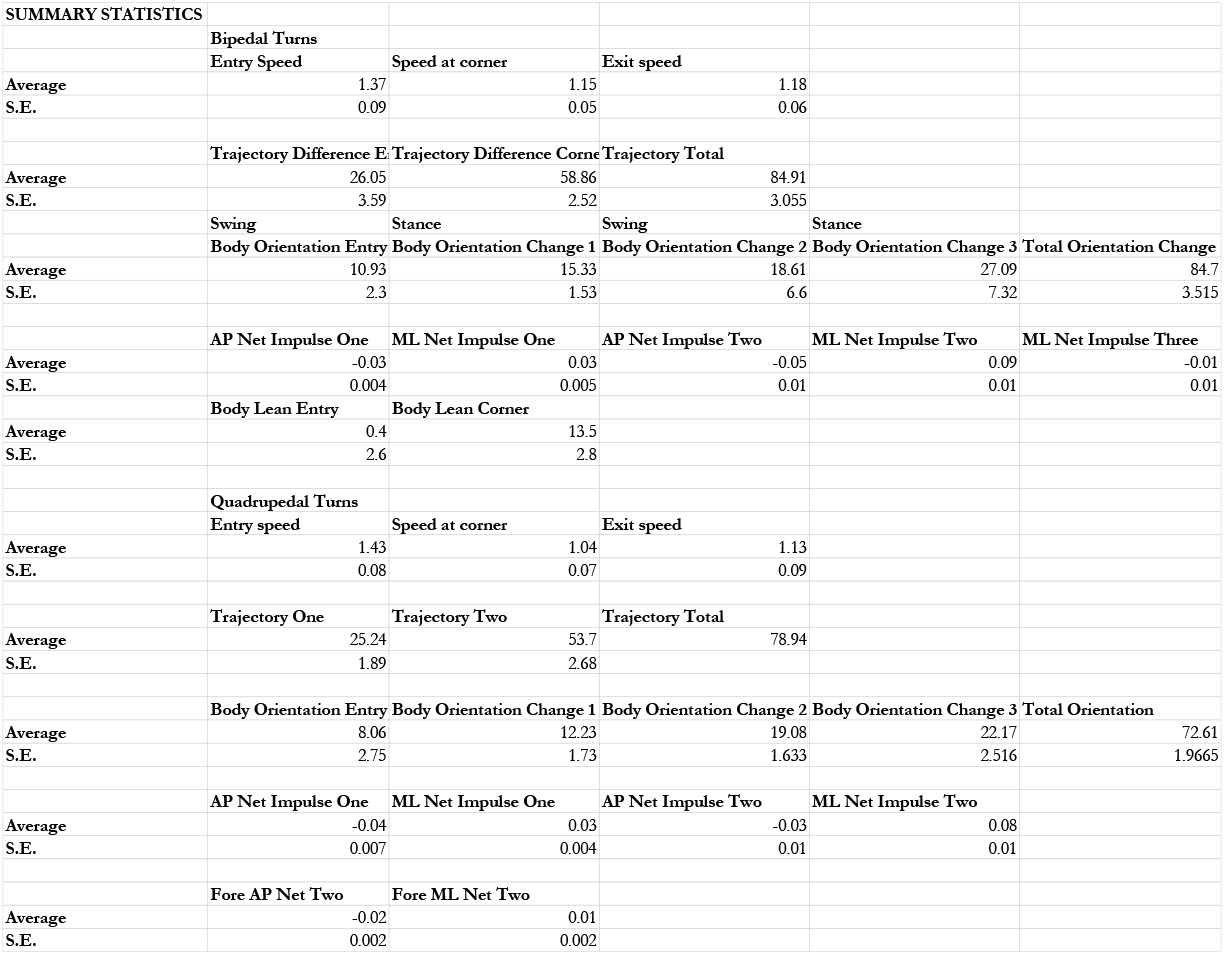
Summary statistics for all included trials. Speeds in ms^-1^. Trajectory and orientation changes in degrees difference of 10 frames before a stance and 10 frames before swing.

Orientation closely mirrored trajectory change leading up to the corner and up to 66% of the 90° trajectory and orientation change was executed on the force plate before entering the corner of the runway (Figure 3). At the corner, orientation did not mirror trajectory change, and much less change occurred (Figure 3). Individuals leaned their bodies from approximately 40° on the left to 40° on the right and body lean corresponded with change in trajectory and ML net impulse (**ρ**<0.05, R^2^=0.45) at the corner (**ρ**<0.03, R^2^=0.46) however, body lean did not correspond with trajectory change at entry, or speed (Figure 4).

**Figure 3:**
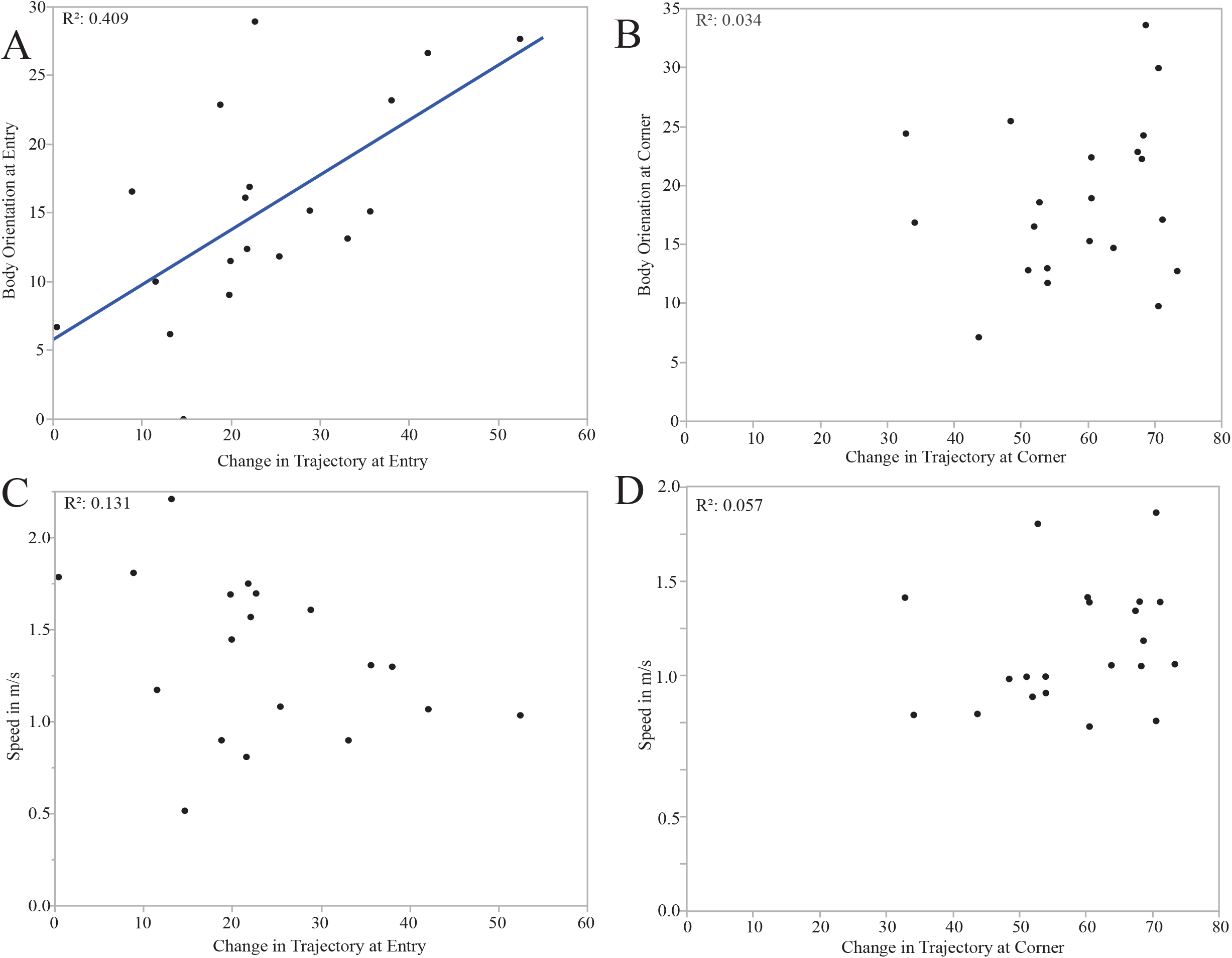
Speed, trajectory changes, and orientation changes at the entry and corner of the turning runway. (A, top left) At entry, body orientation corresponds to trajectory change. (B, top right) At the corner, body orientation does not correspond to trajectory change. (C, bottom left) Speed does not correspond to trajectory change at entry. (D, bottom right) Speed does not correspond to trajectory change at the corner.

**Figure 4:**
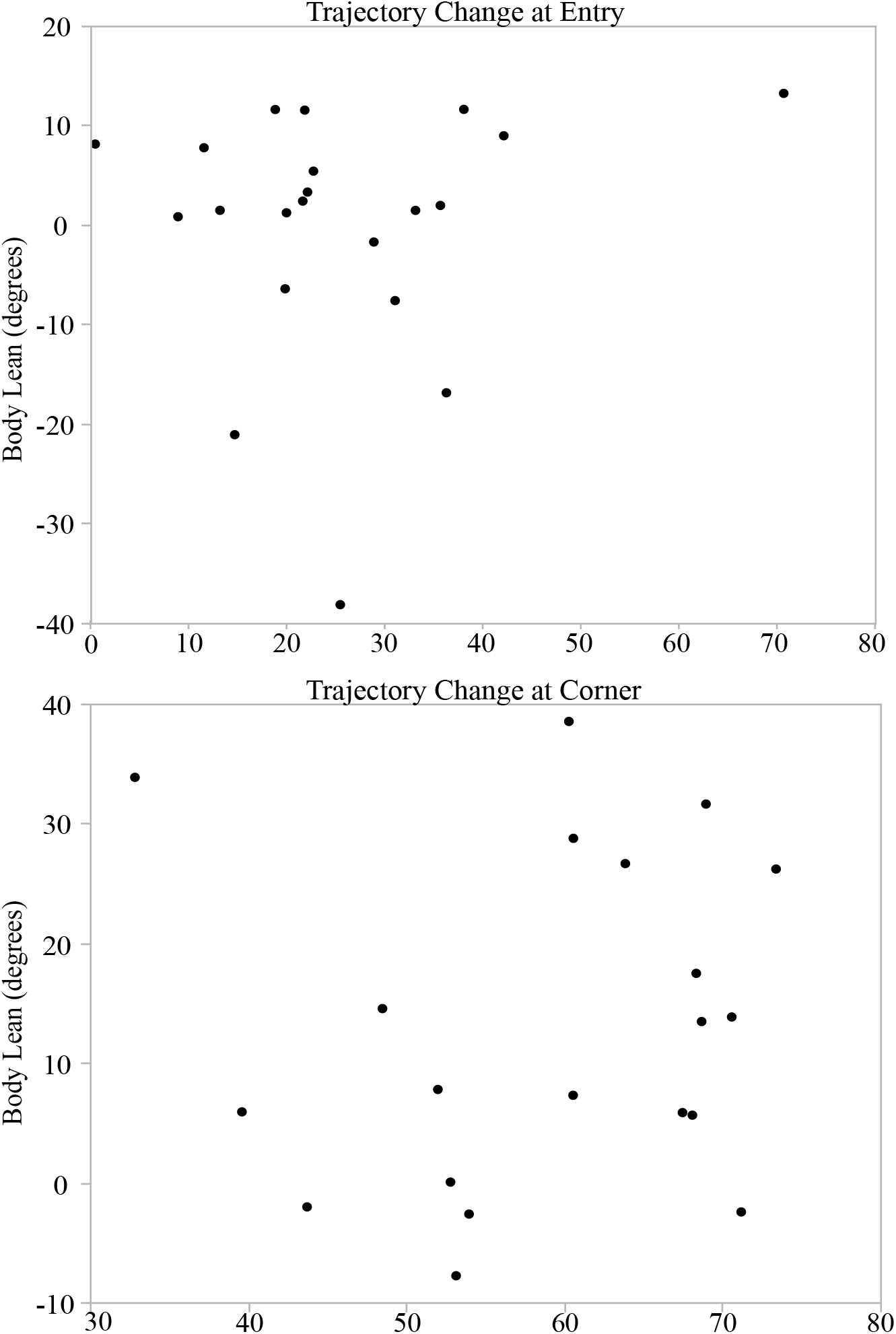
Body lean does not correspond with changes in trajectory. (A) Body lean at turn entry and (B) Body lean at turn corner.

The magnitude of the first trajectory change corresponded with ML forces (**ρ**<0.02, R^2^=0.53), but not AP forces (Figure 5) (ρ>0.05, R^2^: 0.12). The magnitude of the second trajectory change correlated with AP forces (**ρ**<0.01, R^2^=0.45), but not ML forces (ρ>0.05, R^2^=-0.02) (Figures 5). The timing of impulses recorded on each force plate indicate animals applied a braking impulse, followed by a turning impulse (Figure 6). AP forces were applied the ground first, followed by ML forces on the force plate leading up to the corner (entry), as well as on the force plate located in the corner of the runway (Figure 6). AP force peaked at approximately 30% of stance and ML force peaked at 60% of stance at the entry and on the corner (Figure 6). Speed did not significantly correlate with changes in trajectory or orientation (Figure 3) (**ρ**>0.05 in all comparisons).

**Figure 5:**
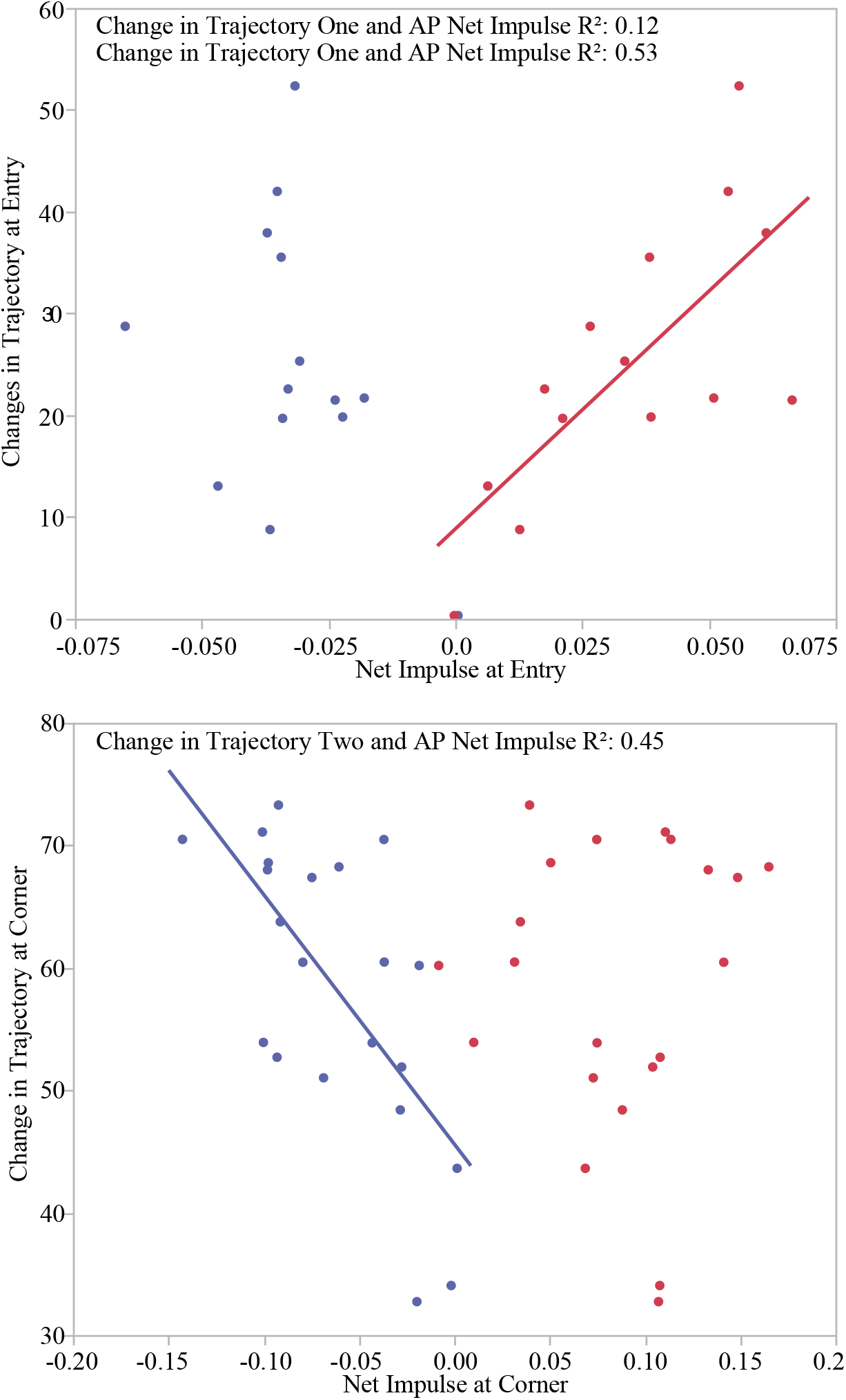
The relationship between AP and ML impulses at the entry (A) and corner (B) of the turning runway. AP (blue) and ML (red).

**Figure 6:**
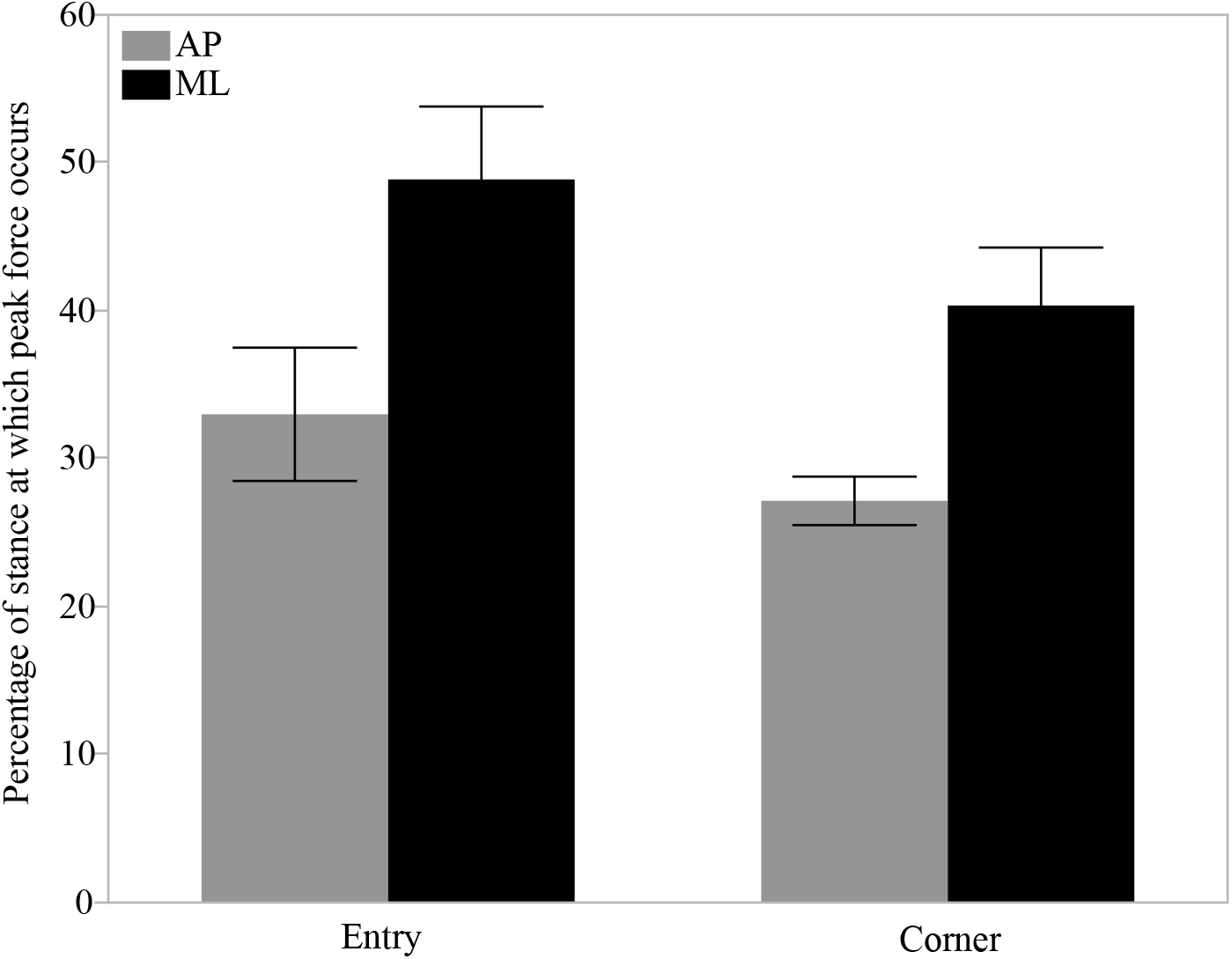
Average (+/1 1 SE) AP and ML forces peak at different percentages of stance at the entry and corner of the turn.

## DISUCUSSION

Critical to determining successful habitat use, competition, and predator-prey interactions in many animals is locomotion. In terrestrial systems, vertebrates transfer muscular forces to Earth’s surface to propel themselves in a desired direction. Therefore, ground reaction forces inextricably link moving vertebrates with their terrain. How forces are applied to the ground affects the intensity of locomotion, thereby affecting the relationship between an individual and its environment, the cost of movement, and determining the outcome of a locomotor behavior. We examined the timing and coordination of forces, trajectory changes, and orientation changes during turning maneuvers by *D. deserti.* Turning is an especially important behavior as it likely increases the probability of surviving a predation attempt (Wilson *et al.* 2013). Specifically, we tested the hypothesis that *D. deserti* change their trajectory by applying a braking impulse and then a lateral impulse. Second, we asked how changes in orientation are achieved, predicting that they closely track changes in trajectory. Data from our experiment indicate braking forces are applied before mediolateral forces, thereby reducing the magnitude of ML forces required to change trajectory (Jindrich *et al.* 2006). Reducing ML forces in turn reduces the likelihood of toppling; thus, braking before the turn means safety is prioritized over turn speed in our study. Initial entrance speeds were not achieved after the corner in this experiment. If our results correspond to wild behaviors, then turns require braking, mediolateral, and accelerative strides. Because turning by kangaroo rats involves braking, turning, and then accelerating in successive order, turning to escape a predator or avoid a competitor likely involves a cost-benefit decision. In some cases, combining a turning maneuver with changes in speed may entail success, while maintaining a steady, straight path is better (Moore *et al* 2015; Wilson *et al.* 2015; Wilson *et al.* 2015).

The individuals in our study adjust their orientation in conjunction with the first change in trajectory. This is especially important as may help visualize any upcoming obstacles, potential refugia, or predators. In any case, the kangaroo rats in our study reorient their bodies before the 90° corner in anticipation of the turn. Natural selection acts on locomotor biomechanics via performance in natural contexts, especially predator-prey interactions (Dickinson et al. 2000, Higham et al. 2016; Wilson et al. 2016). Given our results, changing body orientation early is likely an important behavioral decision during evasive maneuvers (Foster *et al.* 2015). Body lean did not correspond to turning in our study (Alexander 2002). The animals in our study entered and exited the corner with relatively low speeds (1.37 – 1.18 ms^-1^), braking reduced momentum, and their crouched posture yields a low center of mass. Therefore, there was not a large toppling moment.

While animals can only change trajectory while in contact with the ground, reorientation of the body can happen during both stance and aerial phases. For kangaroo rats, reorientation in the air could be facilitated by motions of their long, flexible tails. One advantage of reorienting during the aerial phase would be that the animal would land in position to accelerate along the new trajectory with a parasagittal motion of the legs. Based on observations under natural conditions (Supplementary Video 1), laboratory trials, and digitizing videos, it is likely that kangaroo rats change the relative position and straightness of their tail to facultatively alter rotational inertia (Carrier *et al.* 2001; Walter and Carrier 2002). Future experiments should resolve the role of tails by investigating the role of tail neuromuscular coordination and tail motion in determining the locomotor performance (Jagnandan *et al.* 2014; Jagnandan and Higham 2017).

A vast repertoire of locomotor behaviors enables individuals to acquire food, successfully mate, and escape predators. Whereas straightforward sprint speed has long been shown as key to success in many systems *(e.g.* Husak *et al.* 2006), recent advances point to turning and non-steady maneuvers as important means to catching prey or escaping predators (Moore *et al.* 2015, Wilson *et al.* 2013). We focused on the coordination of turning mechanics under laboratory conditions in *D. deserti.* While we discovered that kangaroo rats initiation trajectory and orientation change before a corner, future work should address how tail kinematics determine how tightly kangaroo rats can turn at varying speeds. In many wild animals, natural and maximal performances are difficult to attain in controlled, laboratory settings. Speeds in our experiment averaged ~ 1.2 ms^-1^ and it was difficult to motivate the animals to hop steadily. Therefore, our study was no different. More work is needed to understand the ranges of speeds and turn angles used in the wild, as well as how terrain mechanics affect turn performance (Young *et al.* 2002).

